# The secreted midbody remnants, MBsomes, as a new class of mRNA nanocarrier applicable to diverse medical fields

**DOI:** 10.1101/2023.09.20.558728

**Authors:** Jae Hyun Kim, Sungjin Park, Ji-Young Seok, Myung-Haing Cho, Soon-Kyung Hwang

**Affiliations:** Department of Veterinary Physiology, College of Veterinary Medicine, Research Institute for Veterinary Science, and BK21 Four Future Veterinary Medicine Leading Education & Research Center, Seoul National University, Seoul, 08826, Republic of Korea; Department of Genetics, University of Wisconsin-Madison, Madison, WI 53706; RNABIO, Inc, Seongnam, Gyeonggi-do 13201, Republic of Korea

**Keywords:** MBsome, mRNA carrier, sucrose cushion ultracentrifugation

## Abstract

The midbodysome (MBsome), a secreted remnant of midbody during cell division, is now known to play a key role in cytokinesis. It was thought that the MBsome is either released into the extracellular space or autophagically degraded by one of its daughter cells. However, recent studies have revealed that MBsomes can be maintained by cells even after cell division is complete, and that they accumulate in the cytoplasm and regulate cell proliferation and survival through integrin and epidermal growth factor receptor-dependent pathways. Here, we examined the ability of MBsomes to act as carriers of mRNAs, a novel function that has not been studied. We found that MBsomes isolated from human lung cancer and stem cells via sucrose cushion ultracentrifugation were 300–400 nm in size and stable for up to 4 days when stored at 4°C. In addition, we confirmed successful expression of the EGFP protein following incubation of the isolated MBsomes with the *EGFP* mRNA at room temperature. These results suggest that MBsomes have the potential to serve as mRNA carriers and therapeutic agents capable of delivering a gene-of-interest.

## 1. INTRODUCTION

Extracellular vesicles (EVs) are nanoscale vesicles derived from the cellular lipid bilayer. EVs have multiple biological functions, including the delivery of components such as proteins and RNA from donor cells to recipient cells, enabling communication between neighboring and distant cells ^1,2^. Due to their natural carrying ability, nanoscale size, and range of contents, EVs have emerged as attractive drug delivery systems and therapeutic tools ^3–5^. However, the application of EVs as carriers of RNA and proteins is hampered by many limitations such as their potential immunogenicity, toxicity and low delivery efficiency ^6,7^. Moreover, the poor storage stability, low yield, low purity (heterogeneity) and weak targeting of exosomes may further limit their clinical applications^8^.

The midbody (MB), a transient structure that forms on the intercellular bridge at the end of cytokinesis, can undergo internalization to form a double membrane-bound organelle ^9^. This organelle, known as the secreted MBsome remnants (MBsomes), forms during ingression of the cleavage furrow as the central spindle microtubules are compacted and cross-linked within a thin intracellular bridge connecting two daughter cells ^10,11^. It is well established that MBsomes plays a key role in orchestrating cytokinesis by recruiting various mitotic kinases, such as Aurora B and Plk1, which are responsible for mediating abscission during the late stages of cytokinesis ^12,13^. In addition, it has recently become clear that MBsomes play a role in regulating cell growth by engaging in cell-cell interactions ^14–17^. Like other EVs, MBsomes are nanosized vesicles that accumulate in cell culture media and can mediate communication with adjacent cells^18^.

Current knowledge of the functions and components of MBsomes is limited. Therefore, we examined whether MBsomes can be used as carriers capable of delivering RNA. First, we developed a method to isolate MBsomes from the culture medium of human lung cancer cells and human adipose stem cells (hASCs) using sucrose cushion ultracentrifugation. Subsequently, we examined the effects of storage conditions on both the stability of the MBsomes and their ability to take up and translate the *EGFP* mRNA. Our findings suggest that MBsomes can be used as carriers for delivering target mRNA.

## 2. MATERIALS AND METHODS

### 2.1. Cell culture

hASCs were purchased from ATCC (PCS-500-011) and were cultured in low-glucose Dulbecco’s Modified Eagle’s Medium (DMEM; 11885076, Gibco, Hillsboro, TX, USA) containing 10% fetal bovine serum (FBS) and 1% penicillin/streptomycin (P/S). The cells (5 × 10^5^) were seeded into 150 mm culture dishes in 15 ml medium and were cultured for 48 hours at 37°C in a 5% CO2 incubator. Subsequently, conditioned medium was obtained from hASCs at passage 7.

The A549 human non-small lung cancer cell line was cultured in high-glucose DMEM (DMEM-HPA; Capricorn Scientific, Ebsdorferground, Germany) containing 10% FBS and 1% P/S. The cells (5.5 × 10^5^) were seeded into 150 mm culture dishes in 15 ml medium and were cultured for 48 hours at 37°C in a 5% CO2 incubator. Subsequently, conditioned medium was obtained from the cells.

### 2.2. Isolation of MBsomes

The secreted MBsomes were extracted from the culture medium of hASCs and A549 cells via sucrose cushion ultracentrifugation. Specifically, the culture medium was centrifuged for 10 minutes at 300 g and 4°C, followed by 10 minutes at 2,000 g and 4°C to remove apoptotic bodies and cell debris, and then 30 minutes at 10,000 g and 4°C. The supernatant was removed and the pellet containing MBsomes was collected. The pellets were resuspended in 34ml phosphate-buffered saline (PBS) and 4ml 30% Sucrose solution and ultracentrifuged for 90 minutes at 100,000 g and 4°C. The MBsome was resuspended in PBS and ultracentrifuged for 90 minutes at 100,000 g and 4°C. Subsequently, the bottom layer was collected and resuspended in 100 µl sterilized PBS to generate the MBsome samples.

### 2.3. Nanoparticle tracking analysis (NTA)

The particle sizes of isolated MBsomes was determined via NTA using the NanoSight LM10 visualization system (Malvern Panalytical, Malvern, UK). MBsomes derived from hASCs were resuspended in 1 ml PBS, and the diluted sample was injected into the LM10 system (with filtered PBS serving as a control). The NTA conditions were as follows: camera level 14, detection threshold value 5, and measurement time 60 seconds (triplicate measurements).

MBsomes derived from A549 cells were diluted 100-fold in Dulbecco’s phosphate-buffered saline (DPBS) and then injected into the LM10 system. The NTA conditions were as follows: camera level 14, detection threshold value 5, and measurement time 90 seconds (triplicate measurements).

NTA was also used to examine the sizes of MBsomes after storage at -80°C, -20°C, 4°C, or 37°C for 4 or 5 days. MBsomes were diluted 100-fold in DPBS and injected into the LM10 system. The NTA conditions were as follows: camera level 14, detection threshold 5, and measurement time 90 seconds (triplicate measurements).

### 2.4. Transmission electron microscopy (TEM)

The morphology of hASC-derived MBsomes was observed using a Talos L120C transmission electron microscope (Thermo Fisher Scientific, Hillsboro, USA). Briefly, after making a grid with a hydrophilic surface, MBsome samples (4 µl) were blotted for 1.5 seconds at 100% humidity and 4°C. Subsequently, the samples were subjected to plunge-freezing in liquid ethane for vitrification using the Vitrobot Mark IV system (Thermo Fisher Scientific). Images were acquired under a transmission electron microscope at 120 kV. The morphology of A549- derived MBsome was observed using a LIBRA 120 instrument (Carl Zeiss AG, Oberkochen, Germany). The sample (5 µl) was first loaded onto the grid for 1 minute, and then PBS was removed. Subsequently, after staining for 20 seconds with 20 µl of 2% uranyl acetate, the sample was dried and subjected to TEM analysis.

TEM was also used to examine the morphology of MBsomes after storage for 4 or 7 days at - 80°C, -20°C, 4°C, or 37°C. After making a grid with a hydrophilic surface, MBsome samples were blotted at 100% humidity and 4°C for 1.5 seconds. The samples were then subjected to plunge-freezing in liquid ethane for vitrification using the Vitrobot Mark IV system (Thermo Fisher Scientific). Images were acquired via TEM at 120 kV.

### 2.5. Immunofluorescence analysis of MKLP1

MBsomes obtained from hASCs and A549 cells were dispensed onto an 8-well chamber slide (30408, SPL Life Sciences, Pocheon-si, Korea) in a total volume of 200 µl PBS. Thereafter, the samples were fixed with 2% formaldehyde for 10 minutes, washed with 200 µl PBS, and then incubated with 0.3% Triton X-100 for 5 minutes. After blocking with 3% bovine serum albumin (BSA) for 2 hours, the samples were reacted with an Alexa Fluor 488-conjugated anti- MKLP1(unique marker of MBsome) antibody (Santa Cruz Biotechnology, Dallas, TX, USA; diluted 1:50 in PBS containing 3% BSA) for 24 hours. Subsequently, the cells were washed with 200 µl PBS, sealed with a fixative solution, and analyzed via confocal laser scanning microscopy (TCS SP5, Leica Microsystems, Wetzlar, Germany).

### 2.6. MBsome stability assay

For stability analyses, isolated MBsomes were stored at 4°C for various periods (1–6 days) and then immunofluorescence analyses were performed. Briefly, MBsome suspended in PBS were seeded onto an 8-well chamber slide (30408, SPL Life Sciences). After 12 hours, PBS in the wells was removed and the MBsomes were fixed in 2% paraformaldehyde for 10 minutes, followed by two washes with fresh PBS. Subsequently, the MBsomes were incubated with an Alexa Fluor 488-conjugated anti-MKLP1 antibody (1:50, diluted in PBS containing 3% BSA) for 12 hours. After washing with PBS, slides were mounted and sealed with mounting medium, and then examined under a confocal laser scanning microscope (TCS SP5, Leica).

### 2.7. Immunofluorescence analysis of EGFP expressed in MBsomes

MBsomes isolated from A549 cells and hASCs were seeded into 8-well chamber slides (30408, SPL Life Sciences), incubated with the CleanCap® *EGFP* mRNA (#L-7601-100; TriLink Biotechnologies, San Diego, CA, USA) for the indicated times, the samples were fixed with 2% formaldehyde for 10 minutes. Also, the samples were washed with 200 µl PBS and then analyzed via confocal microscopy (TCS SP5, Leica). To determine the effect of concentration on *EGFP* mRNA uptake and expression, MBsomes were incubated with various concentrations (2.5, 5, 7.5, 10, and 20 µg/ml) of the CleanCap® *EGFP* mRNA in PBS (total volume: 200 µl) at room temperature for 20 minutes, and EGFP expression was examined by confocal microscopy 24 hours later. To determine the effect of time on mRNA uptake and expression, MBsomes isolated from A549 cells and hASCs were incubated with the CleanCap® *EGFP* mRNA (5µg/ml) in PBS (total volume: 200 µl) at room temperature for 6, 12, 24, 36, or 48 hours and then examined by confocal microscopy.

To determine the effect of storage temperature on EGFP expression, MBsomes were stored at -80°C, -20°C, 4°C, or 37°C for 4 or 5 days, and then incubated with the CleanCap® EGFP mRNA (5µg/ml) at room temperature for 20 minutes. Subsequently, the MBsomes were seeded onto an 8-well chamber slide (30408, SPL Life Sciences) and incubated at 37°C overnight. After fixing with 2% formaldehyde, confocal microscopy was performed.

### 2.8. MBsomes uptake assay

To capacity of MBsomes delivery of target mRNA uptake and expression in cell lines, we used CleanCap® *EGFP* mRNA (5µg/ml) and Pdcd4 mRNA(5µg/ml). MBsomes (1X10^8^ particles/ml) were incubated with 5 µg/ml of the *EGFP* mRNA or Pdcd4 mRNA in PBS (total volume: 200 µl) at room temperature for 20 minutes, and A549 and H460 cells (1X10^4/^wells) were treated with MBsomes Mixtures. After 48h, cells were fixed with 2% formaldehyde for 10 minutes, and then incubated with 0.3% Triton X-100 for 5 minutes. After blocking with 3% bovine serum albumin (BSA) for 2 hours, the samples were reacted with an Alexa Fluor 488- conjugated anti-MKLP1(unique marker of MBsome) or Alexa Fluor 594-conjugated anti- MKLP1 antibodies (Santa Cruz Biotechnology, Dallas, TX, USA; diluted 1:50 in PBS containing 3% BSA) for 24 hours. Subsequently, the cells were washed with 200 µl PBS and MBsomes uptake analyzed via fluorescence microscope STELLARIS8 STED & FALCON, Leica Microsystems, Wetzlar, Germany). To measure of the protein expression level of PDCD4 and GFP in A549 and H460 cells, western blot analysis was performed according to the manufacturer’s instructions. Lysates were mixed with fluorescent master mix and heated at 95°C for 5minutes. Lysates sample, biotinylated Ladder, Primary antibody (PDCD4 antibody; sc- 376430, Santa Cruz Biotechnology, GFP antibody; sc-9996, Santa Cruz Biotechnology), Secondary antibody (Anti-Mouse Secondary HRP antibody), Streptavidin-HRP, chemiluminescence substrate and wash buffer were loaded into the dedicated microplates.

After microplate loading, quantification was performed using the manufacturer’s software (Wes system, ProteinSimple, San Jose, CA).

### 2.9. Statistical analyses

All statistical analyses and were conducted and graphics were generated using GraphPad6 software. The Student’s *t*-test was used to analyze statistical differences. All data represent mean ± standard deviation from at least three individual experiments. A *p*-value < 0.05 was considered statistically significant.

## 3. RESULTS

### 3.1. Characterization of MBsomes derived from hASCs and A549 cells

Figure 1A shows an overview of the experimental procedure used to isolate MBsomes from the conditioned culture medium of hASCs and A549 lung cancer cells. Briefly, the supernatants were centrifuged at low speed to remove cell debris, and then MBsomes were isolated by sucrose cushion ultracentrifugation. The size distribution of the MBsomes was determined by NTA (n = 3). The average size of the A549 cell-derived MBsomes was 310 ± 83 nm, whereas that of the hASC-derived MBsomes was 343 ± 28 nm. (Figure 1B). MBsomes isolated from A549 cells and hASCs displayed a spherical shape with a double membrane (Figure 1C). The red dotted line represents the outer membrane of the MBsome and the yellow circle represents the inner membrane of the MBsome. Fluorescence microscopy also confirmed the expression of MKLP1 in MBsomes from both cell types (Figure 1D). The red and green signals indicate expression of MKLP1 in MBsome isolated A549 cells and hASCs, respectively. Collectively, these results confirmed the successful extraction and purification of MBsomes via sucrose cushion ultracentrifugation.

**FIGURE 1.**
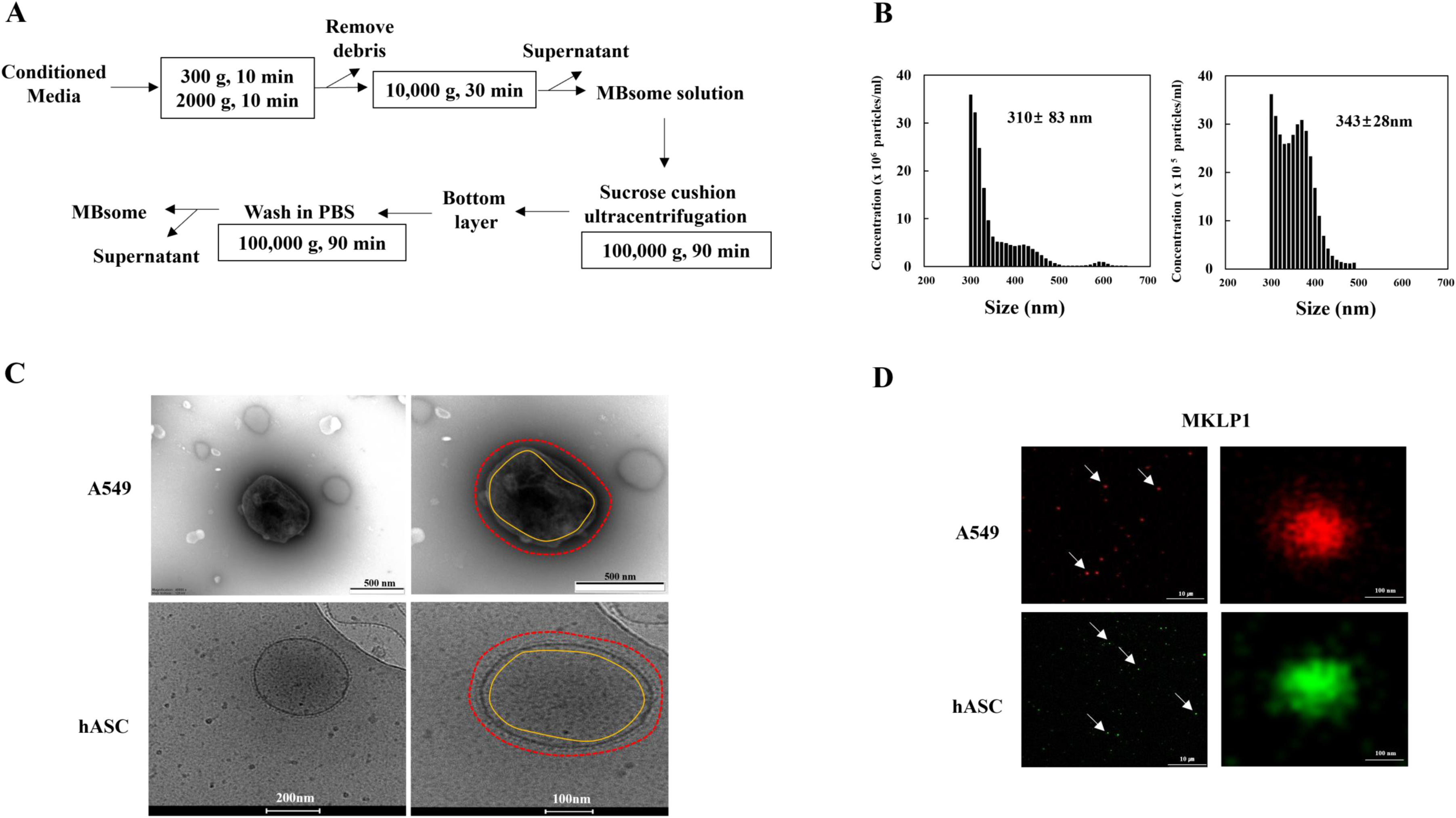
(A) Characterization of MBsomes derived from hASCs and A549 cells. A. Schematic illustration of the method used to isolate MBsomes. (B) NTA showing the size distribution of MBsome derived from A549 cells (left) and hASCs (right). (C) TEM (A549 cells) and cryo-TEM (hASCs) images of isolated MBsomes. D. Immunofluorescence analysis of the MBsomes surface marker MKLP1 in MBsome isolated from A549 cells and hASCs. Arrows indicate MBsomes.

### 3.2. Stability of isolated MBsomes

Several lines of evidence suggest that storage conditions impact the properties of EVs; however, the effects of storage conditions on the properties of MBsomes have not been fully elucidated. To examine their stability, MBsomes isolated from A549 cells and hASCs were stored at 4°C for up to 6 days and then MKLP1 expression was detected by fluorescence immunostaining. MKLP1 expression was decreased significantly after storage at 4°C for 5 or 6 days (Figure 2A and B), indicating that the MBsomes were stable when stored at 4°C for fewer than 5 days.

**FIGURE 2.**
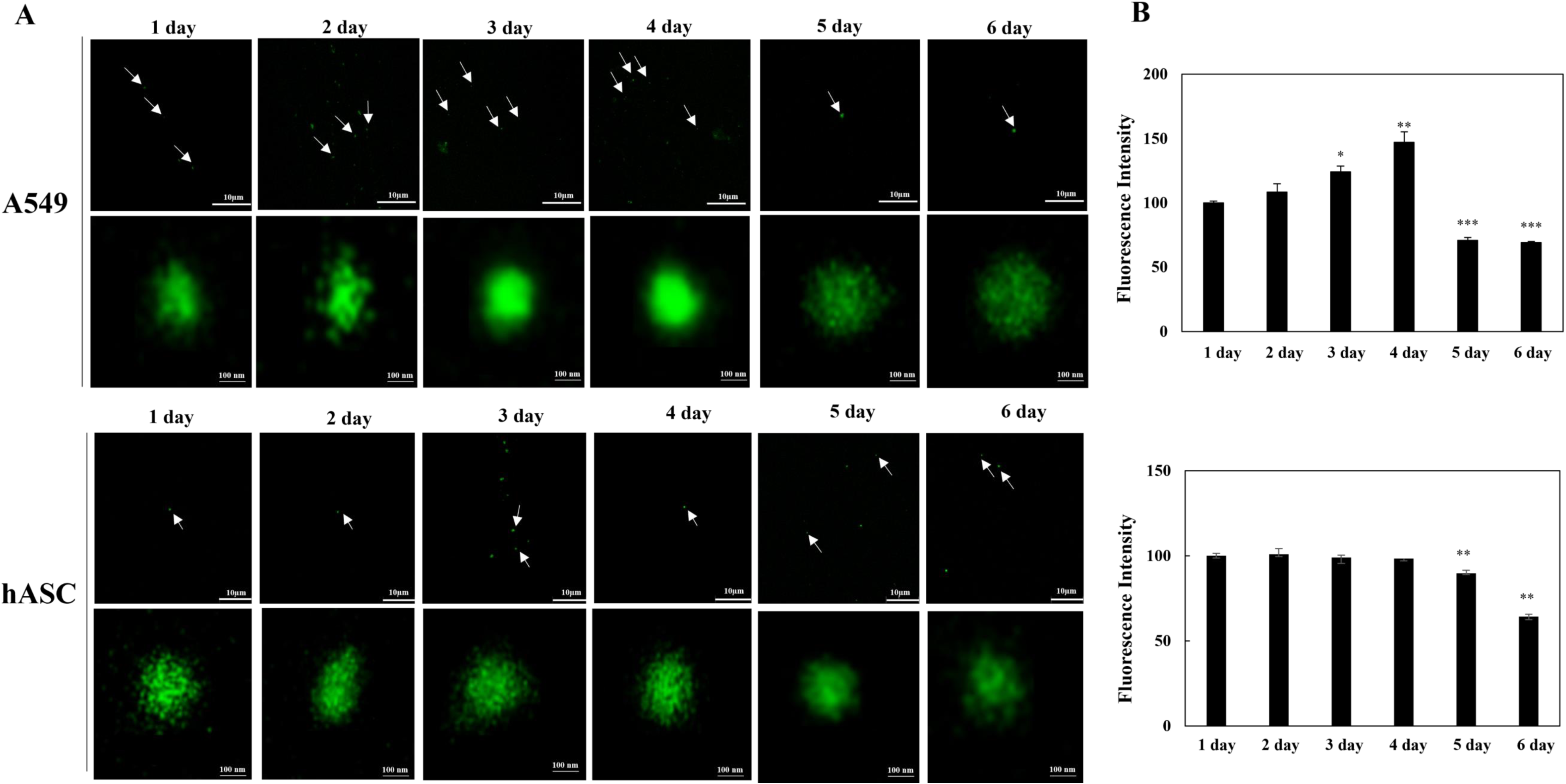
The stability of MBsomes derived from A549 cells and hASCs. (A) and (B) MBsomes isolated from the two cell types were stored at 4°C for 1–6 days and MKLP1 expression was analyzed by immunostaining. Mean ± SD shown for three independent experiments, and statistical significance was calculated by Student’s t-test (*, P<0.05; **, P<0.01; ***, P<0.01).

### 3.3. Function of MBsomes as mRNA carriers

Skop’s group recently demonstrated that MBsomes contain numerous ribosomal proteins, and transcription factors, suggesting that they may function as mRNA translators ^19^. Therefore, we examined whether MBsomes can perform mRNA translation. To this end, MBsomes isolated from A549 cells or hASCs were incubated with various concentrations of the *EGFP* mRNA (2.5, 5, 7.5, 10, and 20 µg/ml) at room temperature for 20 minutes without transfection reagent. Confocal microscopy analysis performed 24 hours later revealed that the EGFP protein was expressed in a concentration-dependent manner (Figure 3A and B). The expression level of EGFP at the 5µg/ml concentration was not lower than those at the higher concentrations; therefore, 5µg/ml was selected as the optimal concentration of mRNA to react with MBsomes. Next, to determine the optimal reaction time for protein expression, the *EGFP* mRNA (5µg/ml) was incubated with MBsome for 6, 12, 24, 36, or 48 hours. In MBsomes isolated from hASCs, the EGFP fluorescence signal was highest for the 48-hour timepoint. However, in MBsomes isolated from A549 cells, the EGFP signal was highest for the 24-hour timepoint (Figure 3C and D). Overall, these results suggest that MBsomes can function as mRNA transporters, and that a minimum mRNA concentration of 5µg/ml and reaction time of 24 hours were required for optimal performance at room temperature.

**FIGURE 3.**
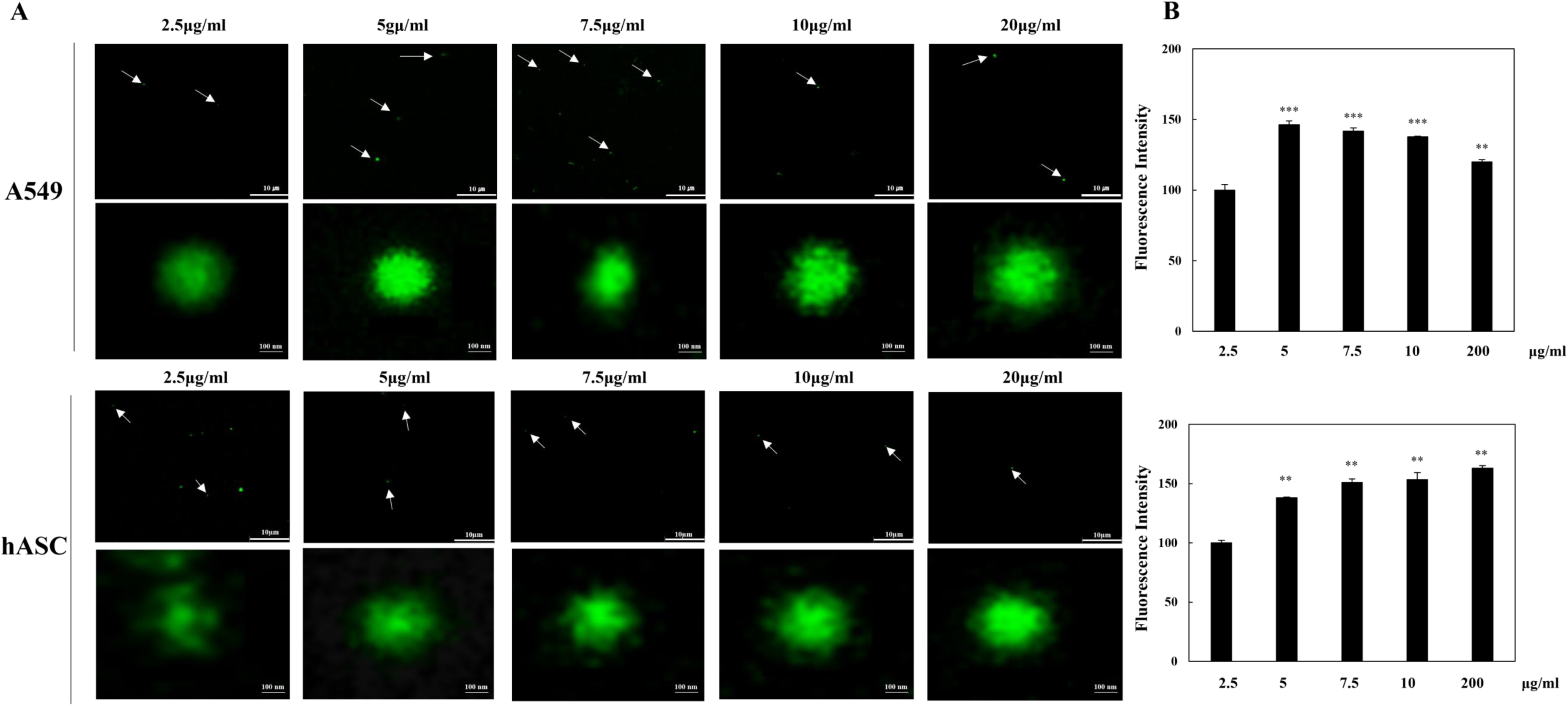
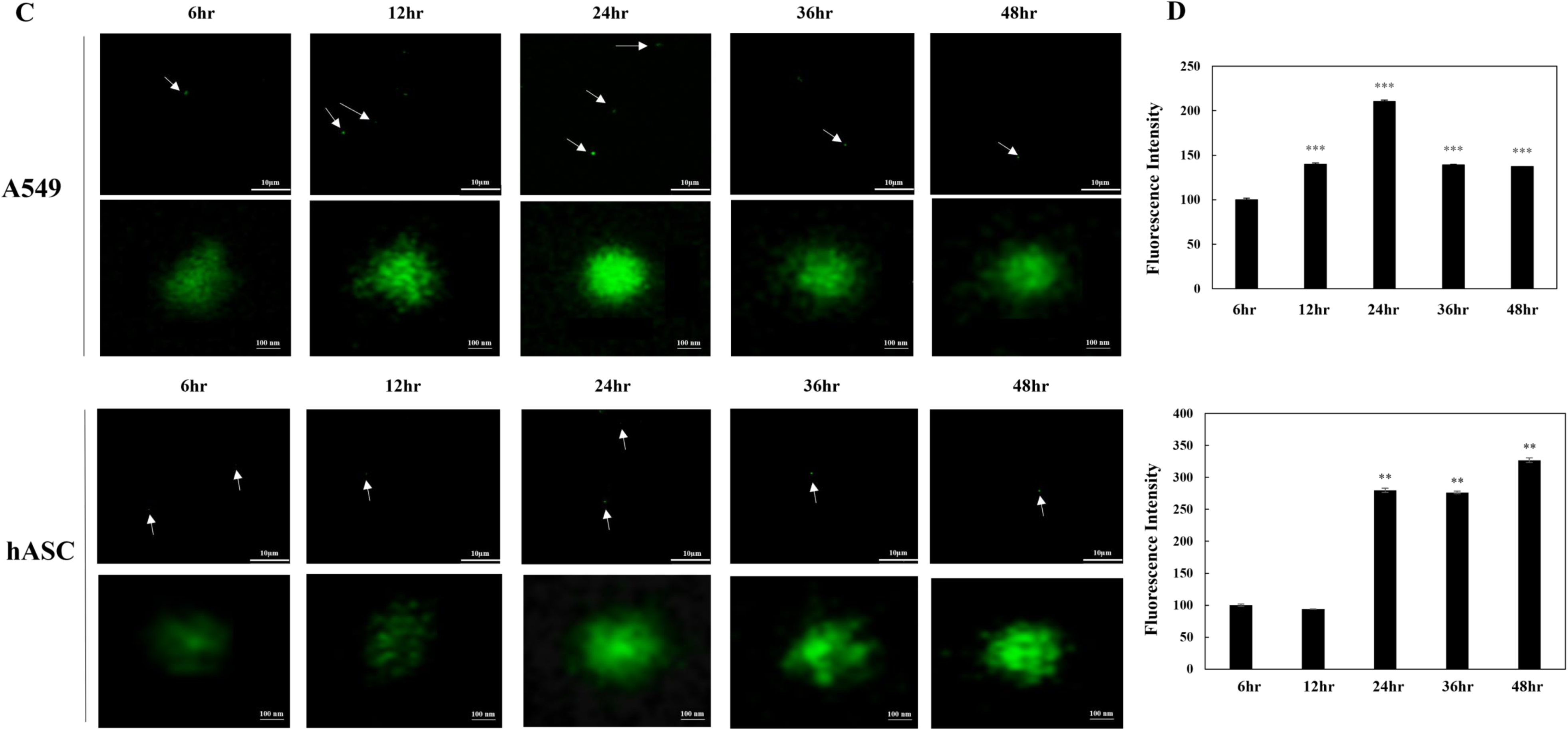
Translation of the *EGFP* mRNA by MBsomes. (A) and (B) MBsomes isolated from A549 cells and hASCs were incubated with the indicated concentrations of the *EGFP* mRNA at room temperature for 20 minutes. EGFP expression was measured by confocal microscopy 24 hours later. Mean ± SD shown for three independent experiments, and statistical significance was calculated by Student’s t-test (*, P<0.05; **, P<0.01; ***, P<0.01). (C) and (D) MBsomes isolated from A549 cells and hASCs were incubated with the *EGFP* mRNA (5µg/ml) for up to 48 hours and EGFP expression was detected by confocal microscopy. Mean ± SD shown for three independent experiments, and statistical significance was calculated by Student’s t-test (*, P<0.05; **, P<0.01; ***, P<0.01).

### 3.4. Optimal storage conditions to maintain the stability and mRNA carrier function of MBsomes

Next, we examined the effects of storage temperature on the stability and function of MBsomes as mRNA transporters. MBsomes isolated from hASCs were stored for 4 or 5 days at -80°C, - 20°C, 4°C or 37°C, and then incubated with the *EGFP* mRNA (5µg/ml) at room temperature for 20 minutes. EGFP expression was detected 24 hours later via fluorescence microscopy. An EGFP fluorescent signal was detected in MBsomes stored under each condition (Figure 4A and B). Cryo-TEM imaging revealed that the shape of the MBsomes after incubation with the *EGFP* mRNA was maintained after the MBsomes were stored for 4 days at all tested temperatures (Figure 4C).

**FIGURE 4.**
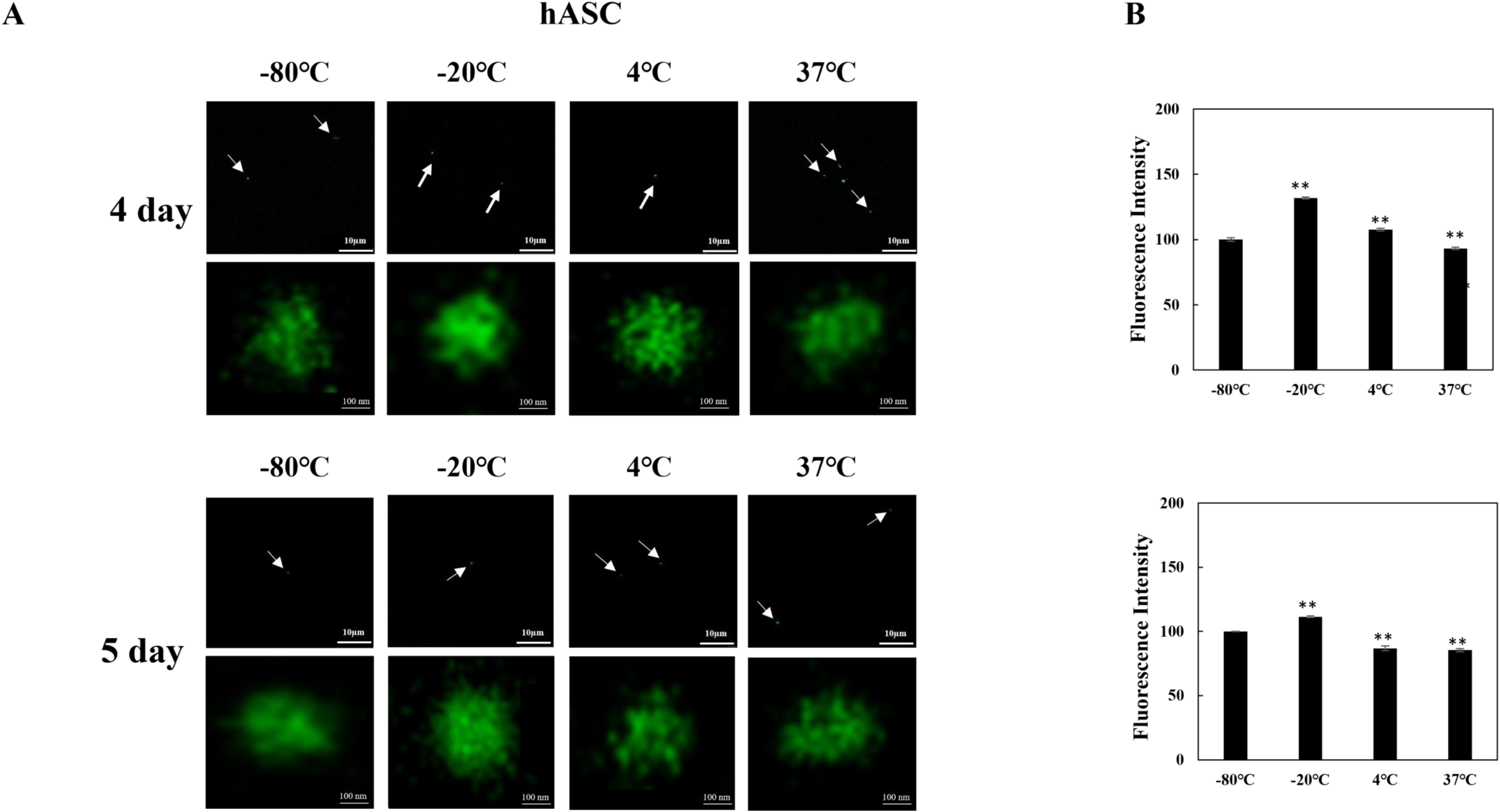
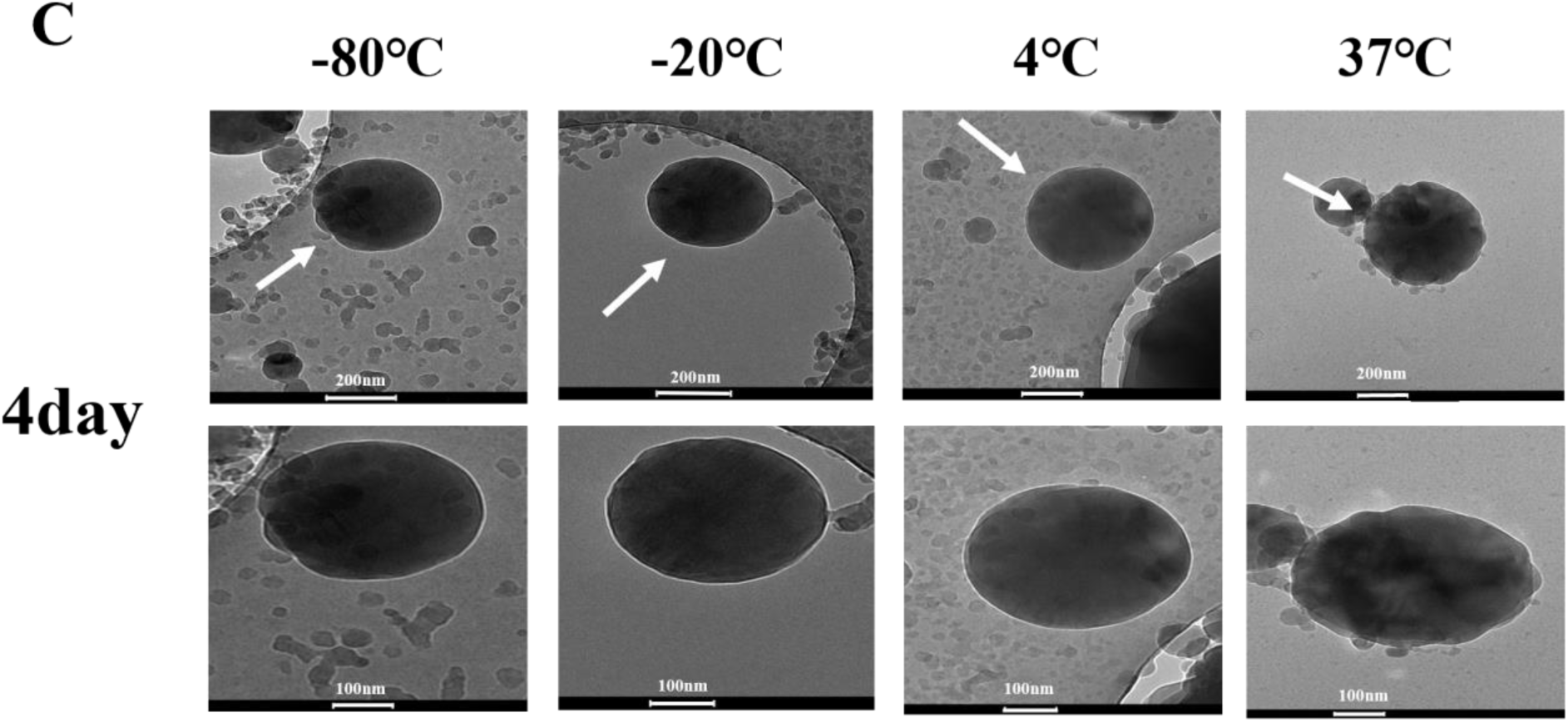
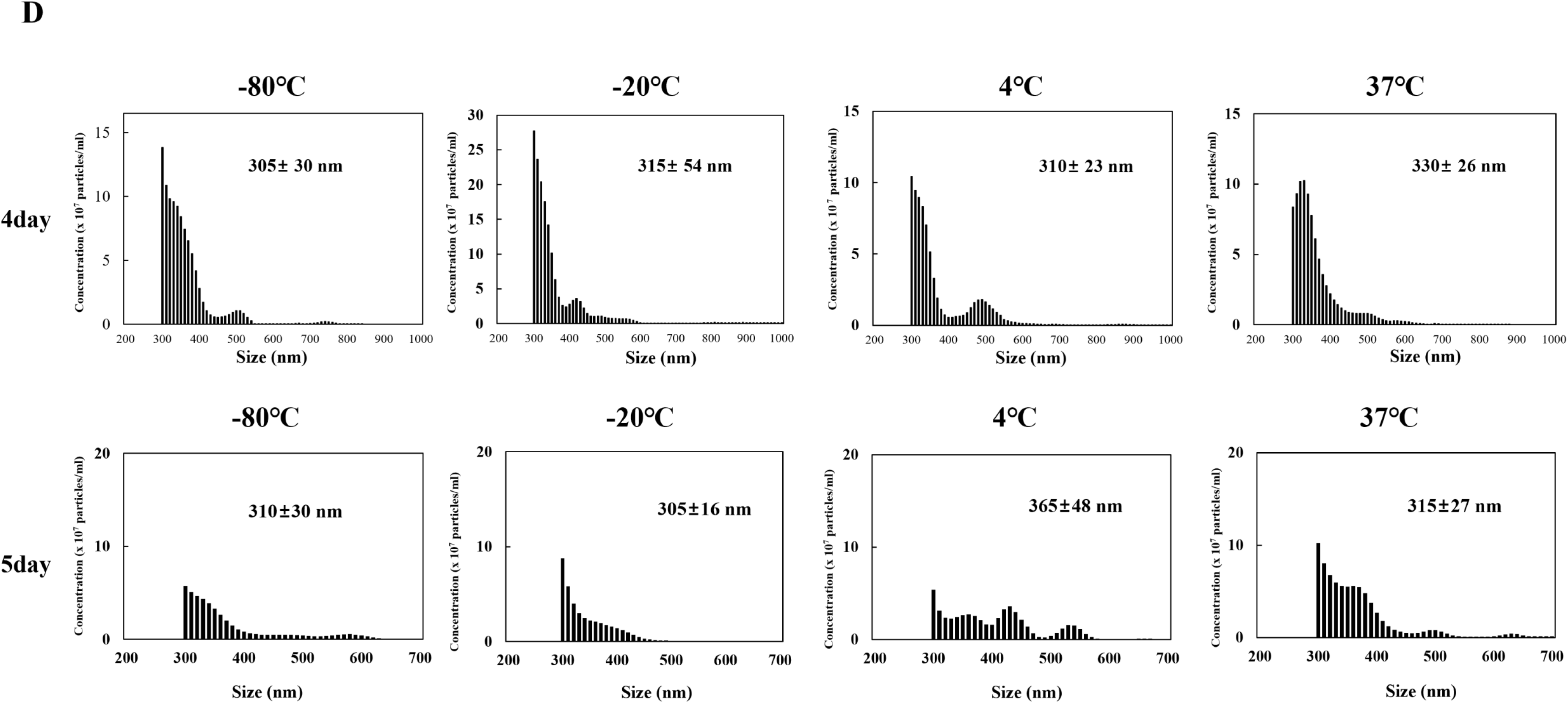
Optimal storage conditions to maintain the stability and mRNA carrier function of MBsomes. (A) and (B) MBsomes isolated from hASCs were stored at the indicated temperatures for 4 or 5 days, and then incubated with the *EGFP* mRNA at room temperature for 20 minutes. EGFP expression was confirmed 24 hours later by fluorescence microscopy. Mean ± SD shown for three independent experiments, and statistical significance was calculated by Student’s t-test (*, P<0.05; **, P<0.01). (C) Cryo-TEM analyses of the morphology of MBsome stored at the indicated temperatures for 4 days. (D) NTA to examine the size distribution of MBsome after storage at the indicated temperatures for 4 or 5days.

In addition, NTA confirmed that the average size of the MBsomes was comparable (approximately 300 nm) after storage at each temperature (Figure 4D). Overall, these results confirmed that the stability and function of MBsomes as a mRNA transporter were maintained after storage for 4 days at various temperatures. Notably, the stability of general EVs stored at 4°C is much lower than that of EVs stored at -80°C ^20^. In contrast with EVs, our findings demonstrate that MBsomes can maintain stability at 4°C. These results strongly suggest that the function of MBsomes as a carrier of a gene-of-interest is maintained upon storage at 4°C for up to 4 days, thereby avoiding the need for storage at -80°C.

### 3.5. *In vitro* uptake of mRNA-loaded MBsome into lung cancer cells

To investigate whether MBsomes derived from human adipose stem cells can act as mRNA carrier by cancer cells, we treated A549 and H460 cells with MBsomes Mixture (MBsomes incubated with *EGFP* mRNA) for 48h, followed by MKLP1 antibody was used to evaluate MBsomes uptake.

Compared to untreated cancer cells, purified MBsomes treated cancer cells showed an increase in MKLP1-positive MBsomes accumulation (red signal). Additionally, we investigated whether MBsomes can deliver mRNA to recipient cells. First, MBsomes isolated from hASCs were incubated with *EGFP* mRNA (5µg/ml) for 20 minutes at room temperature and treated with A549 and H460 cells. Fluorescence microscopy analysis performed after 48h showed that GFP (green) was co-expressed MKLP1(red) in A549 and H460 cells (Figure 5A and B). We then tested the ability of MBsomes to efficiently deliver target mRNAs. When Pdcd4 mRNA was delivered to A549 and H460 cells via MBsomes uptake, PDCD4 was also expressed on MKLP1-positive MBsomes (Figure 5C and D). Fluorescence microscopy revealed that MBsomes can be taken up by lung cancer cells and deliver mRNA. In addition, we investigated whether MBsomes could perform mRNA translation in A549 and H460 cells and verified the protein expression of GFP and PDCD4 through Western blotting. Treatment with lung caner cells treated with MBsomes Mixture (MBsome+ *EGFP* mRNA or MBsome+ Pdcd4 mRNA) significantly increased GFP and PDCD expression compared to untreated cells (Figure 5E and F).

**Figure 5.**
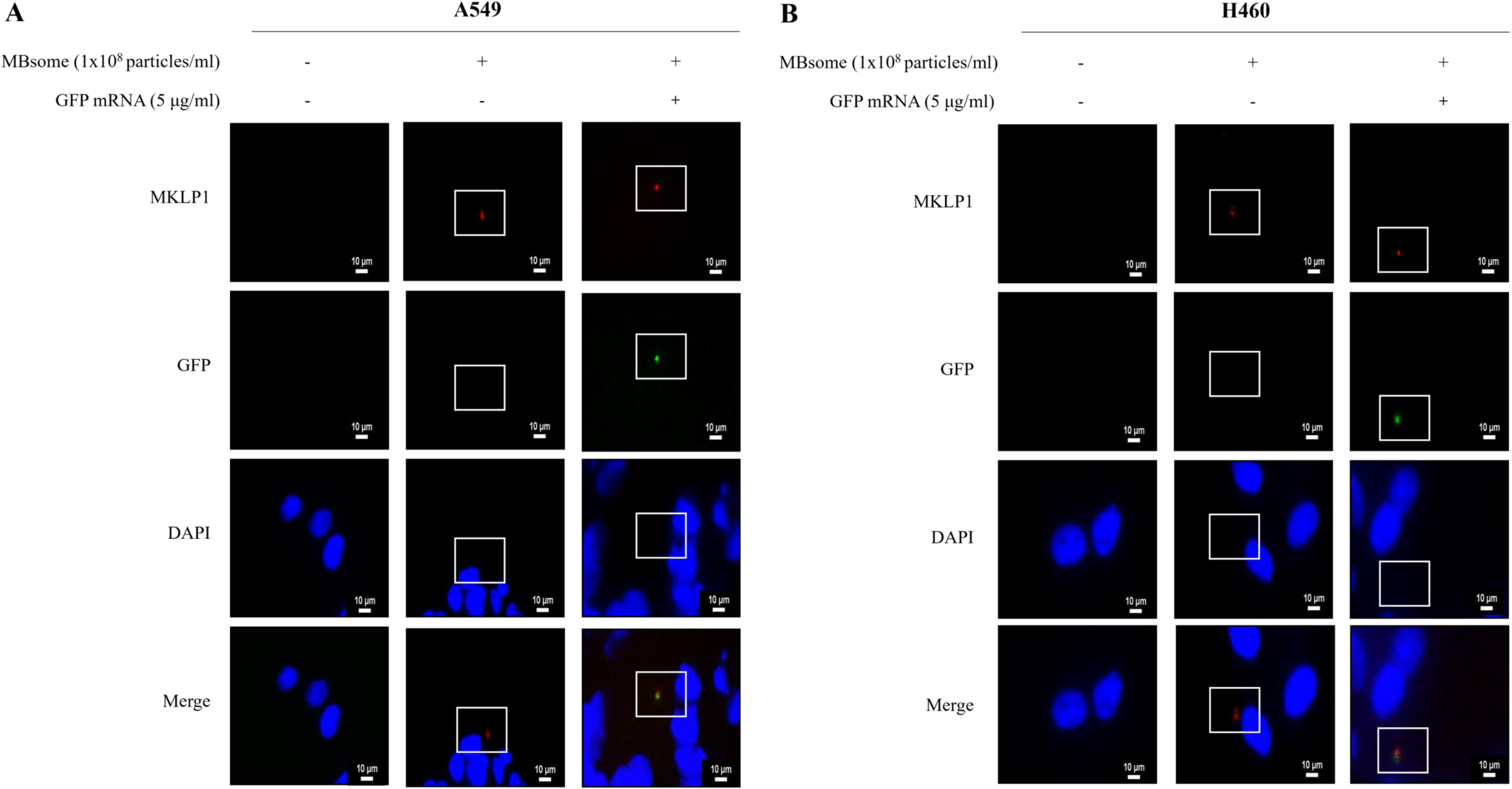
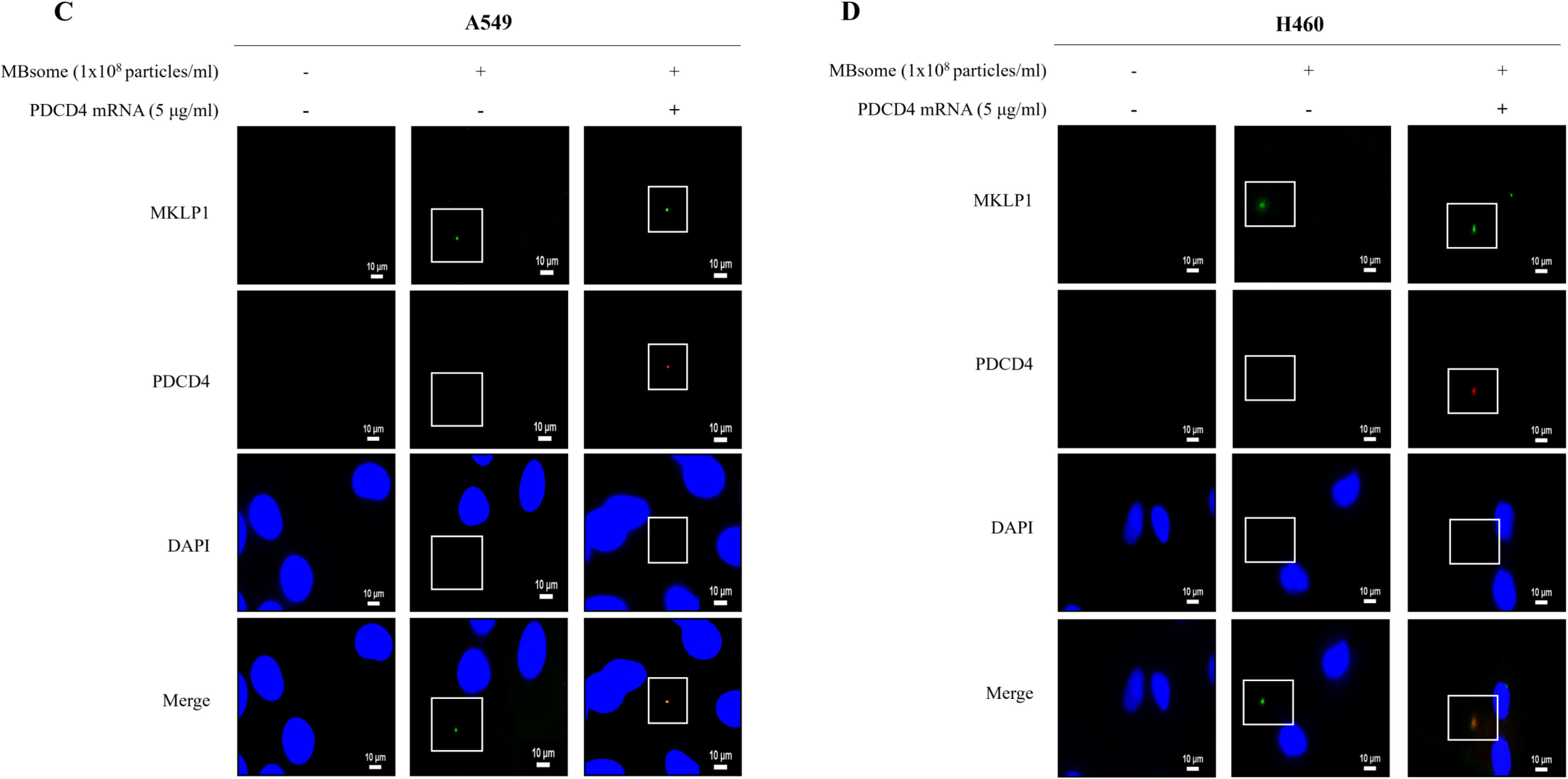
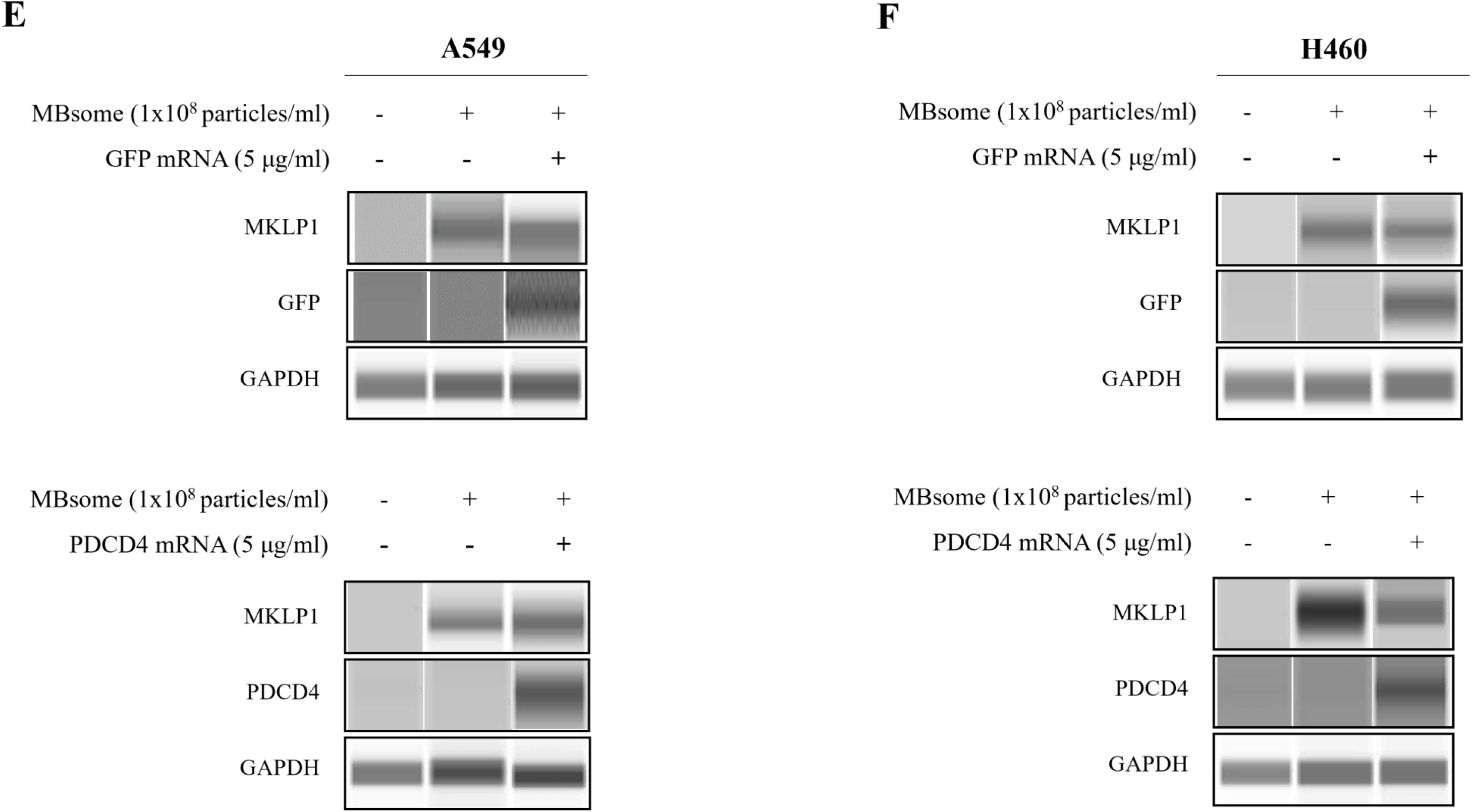
Cellular uptake of mRNA loaded-MBsomes into human lung cancer cells. (A) and (B) cellular uptake of unloaded MBsomes (1X10^8^ MBsomes only) and mRNA loaded- MBsomes (1X10^8^ MBsomes+5µg/ml *EGFP* mRNA) for 48h using anti-MKLP1(MBsomes surface marker; red) antibody and Nuclei were stained with Hoechst stain (blue). GFP expression was confirmed by fluorescence microscopy. (C) and (D) Cellular uptake of unloaded MBsomes (1X10^8^ MBsomes only) and mRNA loaded-MBsomes (1X10^8^ MBsomes+5µg/ml Pdcd4 mRNA). PDCD4 expression was confirmed 48 hours later by fluorescence microscopy. Scale bar, 10 µm. (E) and (F) Western blot analysis to detection proteins GFP and PDCD4 in A549 and H460 cells treated with MBsomes alone or mRNA loaded-MBsomes (*EGFP* or Pdcd4 mRNA) for 48h. Equal protein concentrations of whole cell lysates were analyzed by western blots. GAPDH was used as an equal loading control.

To determine MBsomes loading-mRNA uptake capacity, compare with commercially available cell-expressed mRNA reagents. We transfected with *EGFP* mRNA and Pdcd4 mRNA in A549 and H460 cells using lipofectamine™ MessengerMAX™. We found that the GFP expression of intracellular uptake of MBsomes-loaded *EGFP* mRNA was slightly higher than that of A549 and H460 cells in which *EGFP* mRNA was expressed by lipofectamine transfection reagent (Figure S1). In addition, our results confirmed that MBsomes loaded with Pdcd4 mRNA significantly increased PDCD4 protein expression compared to lipofectamine reacting with Pdcd4 mRNA in H460 cells (Figure S2). These results suggest that MBsomes is suitable for use as an mRNA delivery vehicle.

## 4. DISUCSSION

Sucrose gradient ultracentrifugation has been used widely to isolate conventional EVs ^21 22^. However, it has the disadvantages of low purity and low yield because particles can be broken and impurities are not easily removed ^23–25^. To overcome this problem, we isolated MBsomes by various methods (data not included). We found that 30% sucrose cushion ultracentrifugation is the optimal method to isolate MBsomes.

Here, we used this method to isolate MBsomes, which form at the end of cytokinesis, from the culture medium of two cell types. The characteristics of the isolated MBsomes were examined by NTA, TEM and immunofluorescence analyses. The sizes of MBsomes isolated from lung cancer cells differed slightly from those of MBsomes isolated from hASCs, but both were nano-sized (approximately 300–400 nm). The isolated MBsomes were spherical in shape with a double membrane.

Nano-sized EVs secreted from cells play an important role in intercellular communication ^26 27,28^. Therefore, numerous studies are currently in progress to develop drug and gene delivery systems using EVs ^29–32^. However, natural exosomes may have limitations such as weak targeting and a susceptibility to be quickly cleared from in the body, resulting in poor treatment effects. Therefore, they usually modified to form engineered exosomes for loading with a therapeutic agent or gene^33^. In several studies, drugs or genes were loaded into exosomes via electroporation, chemical transfection, ultrasonic treatment, or extrusion^34 35^.

Here, we investigated whether MBsomes could also act as vehicles for delivering drugs or genes. We found that the EGFP protein was well expressed when MBsomes were incubated with the *EGFP* mRNA without electrical stimulation, outer membrane modifications, or chemical drugs used to deliver genes to other exosomes. These results suggest that MBsomes are a convenient and easy-to-deliver carrier of mRNA.

Exosomes are a promising cell-free therapy, but cannot be stored for a long time ^8^. To protect the biological activity of exosomes and to facilitate their transportation and clinical applications, research of technologies for exosome preservation is ongoing ^36,37^.

In general, EVs retain stability when stored at -80°C and their activity declines sharply when stored at 4°C ^38,39^. A recent study showed that exosomes exhibited the highest stability for 24h when stored at 4 °C, with the highest exosome concentrations and protein levels ^40^. In addition, storage temperature affects the uptake efficiency when EVs are transferred to recipient cells ^41–43^. Here, we found that MBsomes isolated from human lung cancer cells and hASCs were stable for up to 4 days when stored at 4°C, but their stability was reduced on days 5 and 6. These results suggest that MBsomes can be developed as a gene or drug carrier because their stability is maintained at 4 °C for 4 days and they thereby overcoming the limitations of exosomes (short-term storage stability and transport inconvenience). Taken together, these results demonstrate that MBsomes can replace exosomes as a potential off-the-shelf mRNA carrier.

To determine the optimal conditions for MBsomes to act as mRNA carriers, the isolated MBsomes were stored at various temperatures (-80°C, -20°C, 4°C, or 37°C) for 4 or 5 days and then reacted with the *EGFP* mRNA. An EGFP signal was detected in all conditions, and the properties of the MBsomes were not affected by storage temperature. Notably, the EGFP protein was stably expressed when MBsome were stored at 4°C or 37°C for 4 days; therefore, it is likely that stable expression would also be achieved through the storage of MBsome at room temperature for the same period.

## 5. CONCLUSION

Our results strongly suggest that MBsomes can be effectively used as a gene delivery carrier because their stability is maintained at 4°C for up to 4 days; and they easily and conveniently take up target mRNAs.

Moreover, recent data indicate that MBsomes can also be taken up from the intracellular environment, thus potentially serving as platforms for parallel signal transfer between cells. However, further studies will be needed to better understand the regulatory and functional significance of MB inheritance, uptake and accumulation. Here, we have introduced that MBsomes acting as mRNA carriers, can be developed as an off-the-shelf medicine to deliver diverse therapeutic agents and mRNAs for various diseases.

## DATA AVAILABILITY

The data used to support the findings of this study are included within the article.

## AUTHOR’S CONTRIBUTIONS

JHK. SP, MHC, SKH designed the entire study and wrote the manuscript. JHK and SP performed experiment and contributed equally to the data interpretation and the writing and preparation of the manuscript. JYS performed immunofluorescence studies.

## ACKNOWLEDGEMENTS

This research was supported by Basic Science Research Program through the National Research Foundation of Korea (NRF) funded by the Ministry of Science, ICT & Future Planning (NRF-2022R1A2C100780111).

