## Supplementary text for "The secreted midbody remnants, MBsomes, as a new class of mRNA nanocarrier applicable to diverse medical fields"

### METERIALS AND METHODS

##### MBsomes uptake assay

A549 and H460 cells were seeded on culture slide in DMEM medium. MBsomes ($1X10^8$) were incubated with *EGFP* mRNA and Pdcd4 mRNA at room temperature for 20 minutes and then, treated with A549 and H460 cells for 48 hours.

##### *EGFP* and Pdcd4 mRNA transfection

*EGFP* mRNA and Pdcd4 mRNA were mixed with MBsomes or Lipofectamine™ MessengerMAX™ Transfection Reagent (Invitrogen, Waltham, USA) and incubated for 20 minutes at room temperature. After 20 minutes, the mixtures were treated A 549 and H460 cells and used for uptake experiments.

##### Immunofluorescence assay

MBsomes ($1X10^8$) were incubated with 5 µg/ml of the *EGFP* mRNA, Pdcd4 mRNA in PBS (total volume: 200 µl) at room temperature for 20 minutes and treated with A549 and H460 cells ($1X10^4$). After 48h, the samples were reacted with an anti-MKLP1(unique marker of MBsome) antibody, anti-PDCD4 (Santa Cruz Biotechnology, Dallas, TX, USA) for 24 hours. Cellular uptake of MBsomes was analyzed by fluorescence microscope.

##### Westen blot analysis

A549 and H460 cells were seeded in a 60mm dish at $2 \times 10^6$ cells/well in DMEM medium, and treated with MBsome-*EGFP* mRNA mixtures, MBsome-Pdcd4 mRNA mixtures for 48h. GFP and PDCD4 Protein expression levels were detected by Wes system according to the manufacturer's software (Wes system, ProteinSimple, San Jose, CA).

#### FIGURE LEGENDS

##### FIGURE S1

(A) and (B) GFP expression were measured using fluorescence microscopy in A549 and H460 cells after 48h treatment with mRNA loaded-MBsomes ($1X10^8$ MBsomes+5μg/ml *EGFP* mRNA) and transfected *EGFP* mRNA using Lipofectamine™ MessengerMAX™. Scale bar, 10μm.

**FIGURE S2**

(A) and (B) PDCD4 expression in A549 and H460 cells by treatment with mRNA loaded-MBsomes ($1X10^8$ MBsomes+5μg/ml *EGFP* mRNA) and transfected Pdcd4 mRNA for 48h followed by western blot analysis.

### Figure S1.

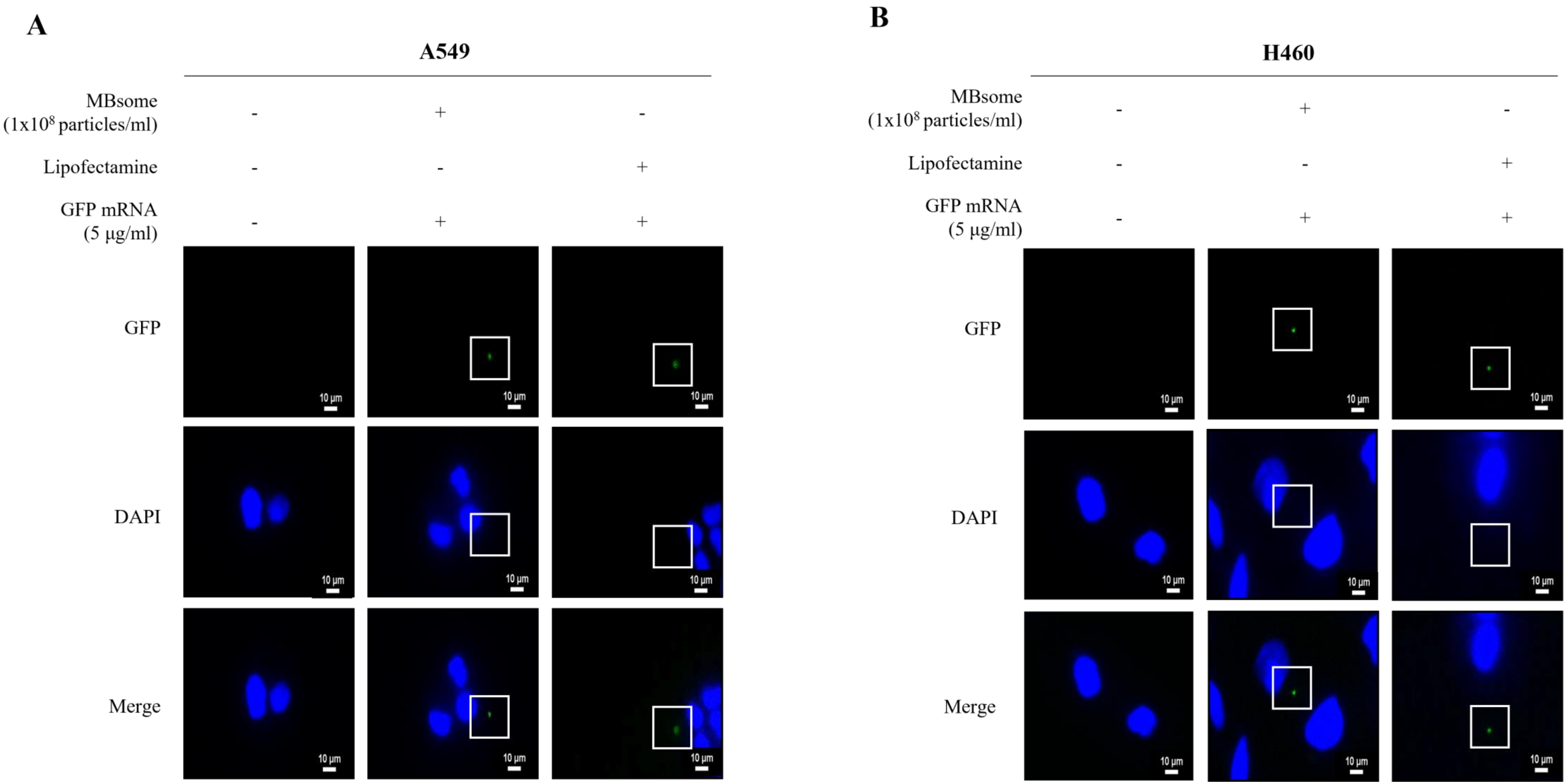
A
A549
MBsome
(1x10⁸ particles/ml)
Lipofectamine
GFP mRNA
(5 μg/ml)
GFP
DAPI
Merge
10 μm
B
H460

### Figure S2.

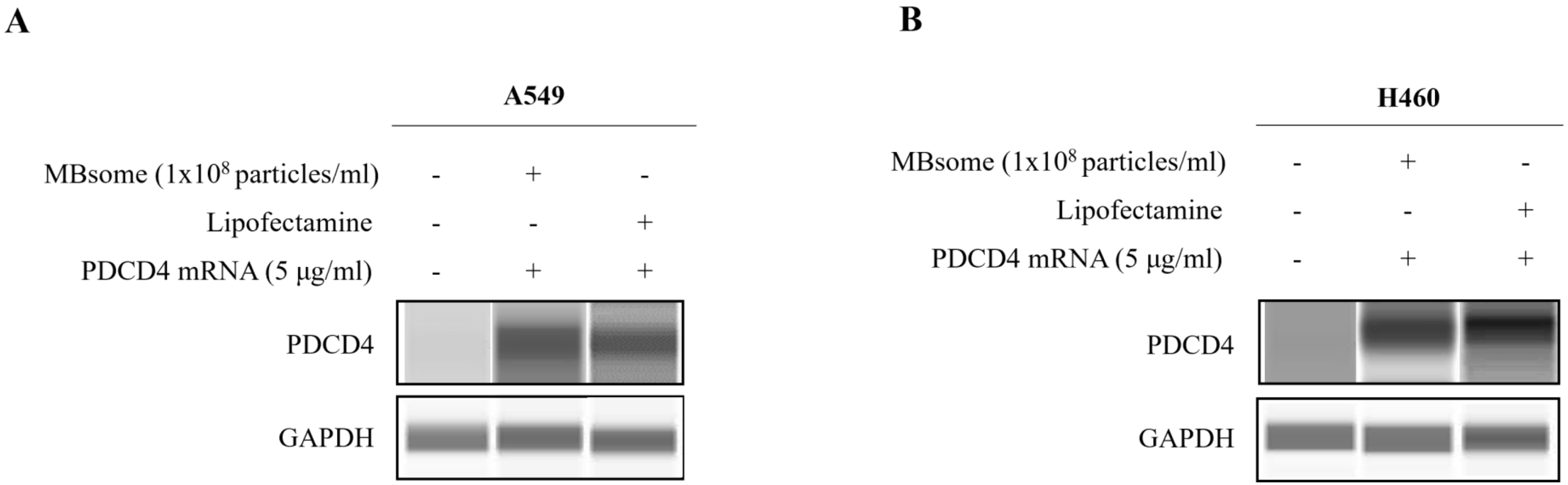
A
A549
MBsome (1x10^8 particles/ml) - + -
Lipofectamine - - +
PDCD4 mRNA (5 μg/ml) - + +
PDCD4
GAPDH
B
H460
MBsome (1x10^8 particles/ml) - + -
Lipofectamine - - +
PDCD4 mRNA (5 μg/ml) - + +
PDCD4
GAPDH
